# Molecular mechanisms of human papillomavirus-related carcinogenesis in head and neck cancer

**DOI:** 10.1101/137653

**Authors:** Farhoud Faraji, Munfarid Zaidi, Carole Fakhry, Daria A. Gaykalova

## Abstract

This review examines the general cellular and molecular underpinnings of human papillomavirus (HPV)-related carcinogenesis in the context of head and neck squamous cell carcinoma (HNSCC) and focuses on HPV-positive oropharyngeal squamous cell carcinoma in areas for which specific data is available. It covers the major pathways dysregulated in HPV-positive HNSCC and the genome-wide changes associated with this disease.

## I. INTRODUCTION

Head and neck squamous cell carcinoma (HNSCC) encompasses a heterogeneous group of malignant neoplasms arising from the non-keratinizing epithelium of the upper aerodigestive tract. Anatomic subsites of HNSCC include the oral cavity, nasopharynx, oropharynx, hypopharynx, and larynx. Squamous cell carcinoma arising from these subsites collectively represents the sixth most common malignancy worldwide, accounting for 932,000 new cases and 379,000 deaths in 2015 [1].

Over the past four decades, striking epidemiological trends have been observed in HNSCC. Although the overall incidence of HNSCC has declined slightly, the relative contribution of each anatomic subsite to the overall incidence of HNSCC has shifted dramatically [2]. Incidence rates of tumors arising from non-oropharyngeal subsites (oral cavity, hypopharynx and larynx) have decreased while the incidence of oropharyngeal squamous cell carcinoma has steadily grown [3]. These subsite-specific epidemiological trends have been attributed to shifts in societal factors that have resulted in changes in exposure to two divergent, but complementary classes of HNSCC risk factors: (1) tobacco and alcohol consumption and (2) human papillomavirus (HPV) infection. Successful public health campaigns in high-income countries are largely credited with achieving population-level decreases in tobacco and alcohol consumption [4] with concomitant declines in tobacco-associated tumors such as non-oropharyngeal HNSCC and lung cancer [5]. Trends toward sexual practices that increase the risk of contracting sexually transmitted pathogens, like HPV, have been linked to the rise in HPV-associated cancers including oropharyngeal HNSCC (OPC) and anal cancers [5,6]. Currently, HPV-positive OPC cases are surpassing the incidence of HPV-positive cervical cancer [3,7].

Human papillomavirus is the most common sexually transmitted infection in the United States and the primary infectious cause of HNSCC [8,9]. Although the spread of high-risk HPV infection is pervasive, the virus is cleared by most people within 18 months [10]. It is believed that persistent infection is necessary for the development of HPV-associated malignancies. The oropharynx exhibits the strongest relationship to HPV. Indeed, HPV-positive oropharyngeal squamous cell carcinoma (OPC), is recognized as a distinct neoplastic entity with a unique molecular, histopathological, epidemiological, and clinical profile [11-13]. Patients with HPV-positive OPC diverge from the classical profile of HNSCC patients in that they are more often younger than 60 years in age and less likely to report a history of heavy tobacco and alcohol consumption [12]. HPV-positive OPC also exhibits marked sensitivity to treatment and a significantly better prognosis than HPV-negative HNSCC [11], observations which have led to the establishment of a staging system specific to this entity [14] and shifts in treatment paradigms [15]. These distinct epidemiological and clinical features are manifestations of the unique underlying biology of HPV-positive OPC. However, much of the research into the role of HPV in HNSCC was conducted before intimate links between HPV and OPC were widely recognized.

## II. HUMAN PAPILLOMAVIRUS

Papillomaviridae is an ancient clade of non-enveloped viruses with a circular double-stranded DNA genome [16]. Within this family, approximately 200 human papillomavirus genotypes, or ‘types’, have been described based on viral genome sequence [17]. Twelve HPV types have been defined by the WHO as high-risk and exhibit high oncogenic potential [18]. At least 10 of these oncogenic HPV types (HPV16, 18, 31, 33, 45, 51, 52, 56, 58, and 59), as well as 6 low-risk HPV types (11, 32, 44, 53, and 81), have been isolated from HNSCC tumors [19-23] (Table 1). HPV16 represents the primary viral cause of HNSCC and is identified in at least 87% of HPV-positive OPC [24]. HPV18 and HPV33, the next most prevalent types, account for most of the remainder of HPV-positive HNSCC [24].

**Table I.** HPV genotypes identified in HNSCC

| High-risk | Low-risk |
| --- | --- |
| Genotype | Genotype |
| 16 | 11 |
| 18 | 32 |
| 31 | 44 |
| 33 | 53 |
| 45 | 57 |
| 51 | 81 |
| 52 |  |
| 56 |  |
| 58 |  |
| 59 |  |

All papillomaviruses possess a genome of approximately 8 kilobases encoding 8 open reading frames (ORFs) that enable viral genome replication and viral particle assembly (Figure 1). ORFs of the HPV genome are organized based on the timing of expression with respect to the viral life cycle: early (E1, E2, E4, E5, E6, and E7) and late (L1 and L2) genes [25]. The E1 viral helicase and E2 DNA-binding protein directly mediate viral genome replication, while E4, E5, E6, and E7 are accessory proteins that coordinate viral genome amplification and virulence. The late genes L1 and L2 encode viral capsid proteins necessary for the final stages of virion assembly and mediate viral entry into future host cells. The functional diversity of this limited set of ORFs is expanded through complex patterns of viral transcript splicing and stage-specific gene expression that is linked to host cell differentiation [26].

**Figure 1:**
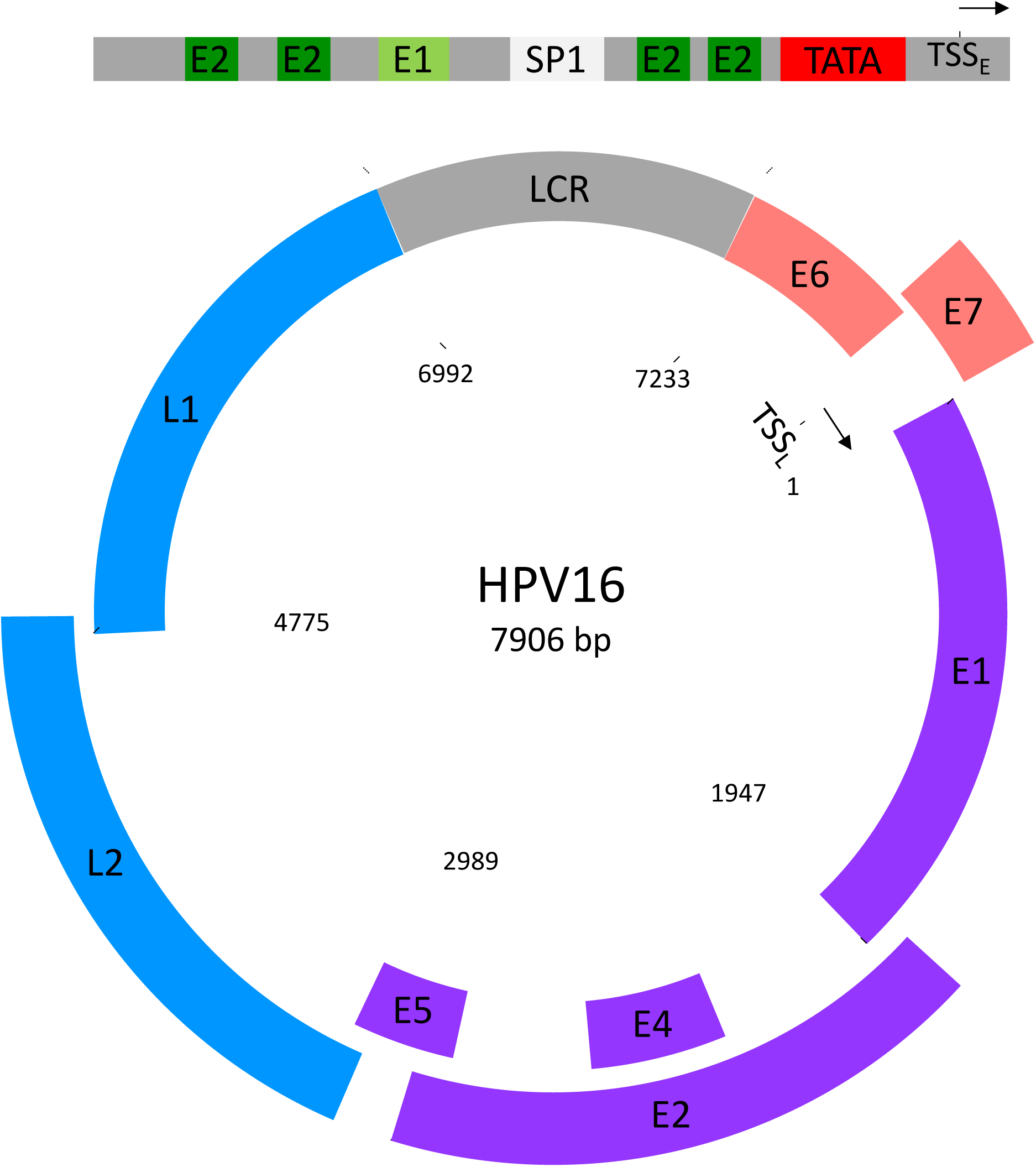
The HPV16 genome. Schematic of HPV16 genome designed from Ref Seq entry: NC_001526. Viral genes E6 and E7 are marked in light red; E1, E2, E4, E5 in purple; and L1 and L2 in blue. Green regions marked E1, E2 are binding sites for the corresponding proteins. LCR: long control region; TSS_E_: transcription start site for viral early genes; TSS_L_: transcription start site for viral late genes; TATA in red represents the TATA box transcription factor binding site. SP1 represents the binding site for the host SP1 transcription factor. Although numerical base markers represent viral genomic positions, the coding elements are not to scale.

### Infection

The host cells for HPV infection are keratinocyte progenitors located in the basal layer of stratified squamous epithelia and adhered to the epithelial basement membrane. Experimental models suggest that infection requires viral access to the basement membrane [27] (Figure 2). In the epidermis and anogenital tract, HPV gains access to basal cells through micro-abrasions that occur during sexual or other direct physical contact [28].

**Figure 2:**
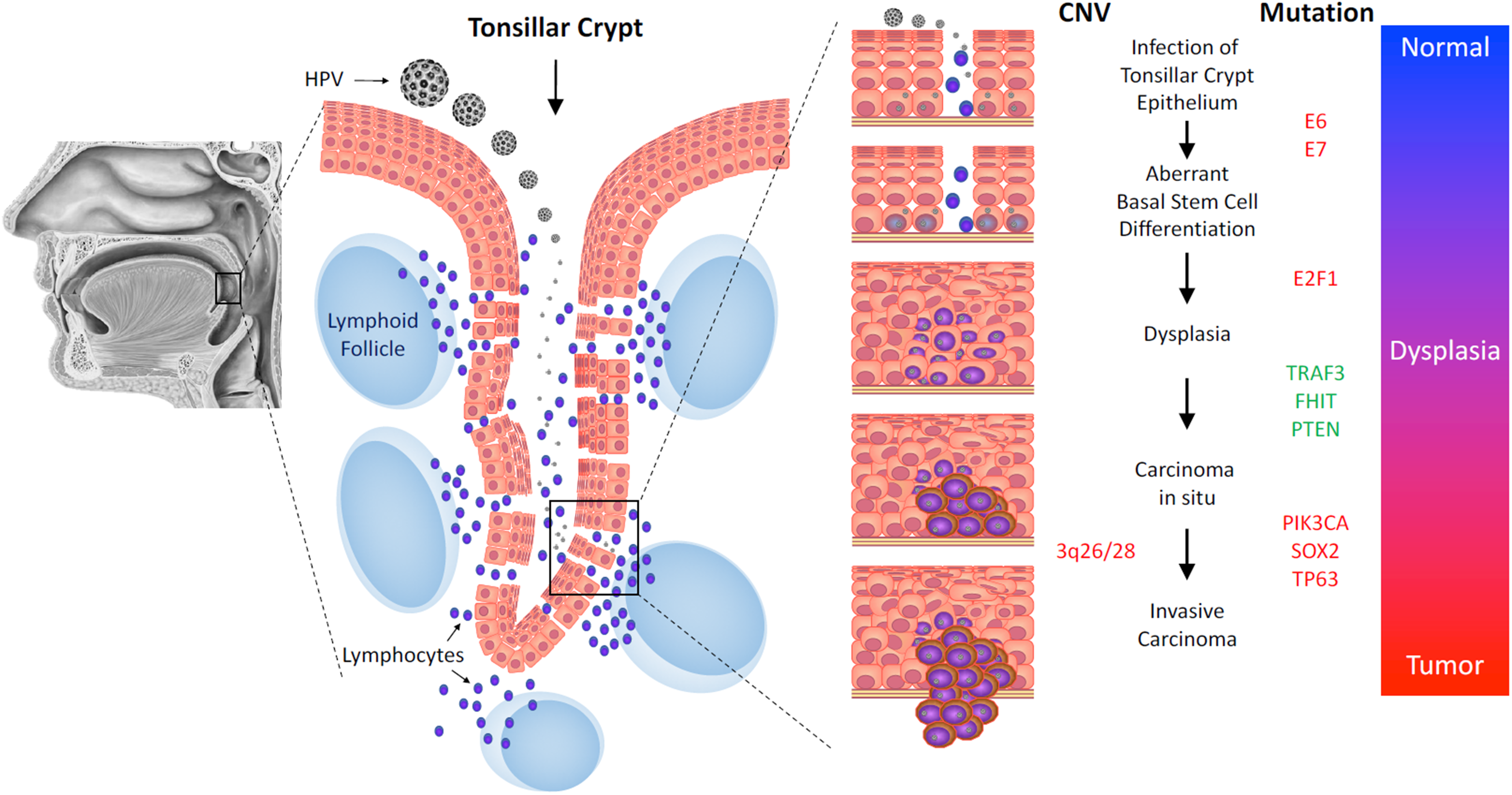
Tumor progression model for HPV-related carcinogenesis. (A) In the head and neck, HPV demonstrates tropism for lymphoid-associated structure of the oropharynx, including the palatine and lingual tonsils. (B) In the oropharynx, HPV gains access to basal keratinocyte progenitors through fenestrations in the reticulated epithelium of the tonsillar crypts. (C) Infection of the tonsillar epithelium result aberrant basal cell differentiation, dysplasia, carcinoma in situ, and finally invasive carcinoma. (D) Hypothetical model for somatic mutations in a multistage tumor progression model. Genes and loci in red are upregulated or activated or show increase in copy number. Genes in green undergo loss of function mutation or deletion.

In the oropharynx, HPV infection may occur in the absence of epithelial abrasion. The palatine, lingual, tubal, and adenoid tonsils are lymphoid structures collectively known as Waldeyer’s tonsillar ring. Constituents of Waldeyer’s ring possess a specialized reticulated squamous epithelium infiltrated with lymphoid tissue. The reticulated epithelium contains a fenestrated, discontinuous basement membrane thought to allow immune cells access to oral antigens [29]. These fenestrations also represent natural interruptions in epithelial barriers that likely provide HPV access to basal keratinocytes in the absence of traumatic epithelial disruption (Figure 2) [30]. Indeed, the unique epithelial properties of Waldeyer’s ring may explain the disproportionate tendency of HPV to cause squamous cell carcinoma of the palatine and lingual tonsils [31].

Upon reaching the basal keratinocyte, HPV preferentially binds components of the extracellular matrix. The L1 capsid protein directly engages basement membrane heparin sulfate proteoglycans [32], triggering conformational changes in L1 and L2 that transfer virion particles to host cellular uptake receptor(s) necessary for viral internalization. Although cellular HPV receptors mediating viral entry have not been definitively identified, accumulating evidence suggests that tetraspanin-enriched microdomains on the plasma membrane serve as the primary platform for viral entry into the cell [33]. Once internalized, virion particles undergo endosomal transport, uncoating, and cellular sorting into the nucleus [34].

Upon entry and uncoating, the circular viral DNA is transported to the nucleus where it exists as an episome, a genetic element separate from the host cell genome, that employs host cell enzymes to replicate its genome along with host chromosomes and is maintained at low-copy number (∼50-100 copies per cell) [35]. Tight regulation of low-copy replication of the viral genome in basal keratinocytes serves as one mechanism by which HPV evades the host immune system. Oncoproteins E6 and E7 are expressed prior to productive viral replication, driving cell cycle entry and cell proliferation in the basal and parabasal cells [26]. In oncogenic HPV genotypes, cell cycle dysregulation by E6 and E7 constitutes the initial steps driving HPV-related carcinogenesis.

The expression of viral genes and virion production increases rapidly in daughter keratinocyte progenitors undergoing differentiation and progressing toward the mucosal surface [36]. In normal stratified squamous epithelia, only basal keratinocyte stem cells possess the potential to proliferate. Asymmetric cell division of these keratinocyte stem cells leads to two daughter cells: one renewed basal stem cell and one cell destined to become a terminally-differentiated keratinocyte. HPV-encoded genes perturb key keratinocyte differentiation and proliferation pathways to favor virus production and viral life cycle completion. In high-risk HPV types, these perturbations predispose cells to neoplastic transformation.

## III. CANONICAL MECHANISMS OF HPV-DRIVEN CARCINOGENESIS

Knowledge derived from the study of HPV-related cervical cancer composes the general framework for understanding mechanisms of carcinogenesis in HPV-related HNSCC. High-risk HPV types encode virulent alleles of two viral proteins, E6 and E7, that endow viral oncogenic potential by targeting molecular pathways that underlie neoplastic transformation. Downstream consequences of high-risk HPV E6 and E7 action includes the promotion of cell cycle progression, disruption of keratinocyte differentiation, keratinocyte immortalization, genomic instability, and neoplastic transformation [37].

A central advance in the understanding of HPV-related carcinogenesis arose with the elucidation of mechanisms for the oncogenic potential of E6. High-risk HPV E6 proteins bind to and induce ubiquitin-dependent proteolysis of the p53 tumor suppressor protein [38]. P53 is the protein product of the *TP53* gene, a regulator of the DNA-damage response [39], G_2_/M cell cycle transition [40], and the most commonly mutated tumor suppressor gene in cancer (Figure 3) [41]. Only E6 alleles from high-risk HPV types bind p53 and the strength of the binding affinity of E6 alleles for p53 correlates with that HPV type’s oncogenic potential. For example, HPV16, which is responsible for more HPV-related HNSCC than HPV18 [42], encodes for an E6 allele that binds p53 with twice the affinity of the HPV18 E6 allele [43].

**Figure 3:**
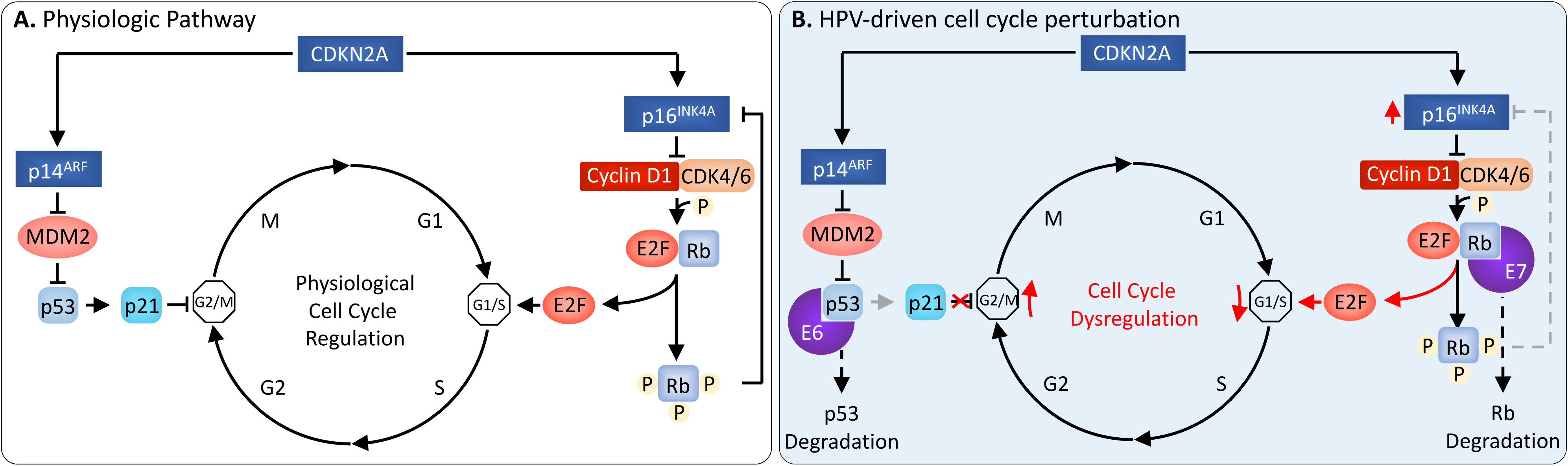
Cell cycle perturbation by human papillomavirus. (A) Schematic representing physiologic cell cycle regulation. Tumor suppressor proteins are marked blue, proto-oncoproteins are marked red. The *CDKN2A* gene encodes p14^ARF^ and p16^INK4A^ proteins. p14^ARF^ inhibits MDM2, thereby disinhibiting p53 to activate p21 and stop progression through the G2/M checkpoint into mitosis. p16^INK4A^ inhibits the CyclinD1/CDK4 and CyclinD1/CDK6 complexes. These complexes catalyze phosphorylation of the retinoblastoma protein (Rb), inducing it to release E2F family transcription factors to enter the nucleus and activate transcription of S-phase promoting transcripts. Phosphorylation of Rb also results in feedback inhibition of p16^INK4A^ expression. (B) HPV E6 oncoprotein binds and targets p53 for degradation, resulting in loss of G2/M checkpoint regulation. HPV E7 oncoprotein binds and targets Rb for degradation, resulting in the nuclear translocation of E2F and promotion of S-phase transition. In addition, downregulation of Rb results loss of feedback inhibition and overexpression of p16^INK4A^.

In an analogous mechanism to the action of E6 on p53, the HPV E7 oncoprotein exerts its oncogenic effect by facilitating the proteolytic destruction of Retinoblastoma (Rb)-family tumor suppressors [44]. Rb enacts its primary inhibitory effect on the cell cycle by regulating the nuclear accumulation of E2F and thereby controls the G1 to S-phase cell cycle transition (Figure 3). E7 induces enhanced degradation of Rb protein via a ubiquitin-proteasome pathway [44]. Degradation of Rb releases E2F family mitogenic transcription factors into the nucleus, promoting entry into S-phase and activating proliferative transcriptional programs [45,46]. Degradation of Rb also releases *CDKN2A* gene expression, which codes for the tumor suppressor p14^ARF^, as well as p16 (or p16^INK4A^ - a surrogate marker of HPV infection [47,48]). Low-risk E7 binds Rb at an affinity that is insufficient to induce levels of Rb degradation necessary to promote neoplastic transformation (Figure 3) [49,50].

Beyond these well-characterized actions, E6 an E7 modulate the activity of numerous other cellular proteins. Recent data suggest that viral oncoproteins E6 and E7 bind and modify the activity of numerous proteins and molecular pathways [51]. E6 has been reported to interact with 83 cellular proteins and E7 with 254 [52].

## IV. GENOME-WIDE SOMATIC MUTATIONS OF HPV-RELATED HNSCC

Notable studies have shed light on the unique mutational profile of HPV-positive HNSCC. They have served to distinguish HPV-positive HNSCC from their HPV-negative counterparts based on molecular, rather than simply clinical, markers. Insight gained from their contributions provide additional targets for which prevention and therapy can be developed.

The Cancer Genome Atlas (TCGA) consortium presented the most comprehensive HNSCC genome-wide analysis to date [1]. Analysis of 279 HNSCC, of which 36 were HPV-positive, identified recurring activating mutations in *PIK3CA* in HPV-positive HNSCC [53] [1]. *PIK3CA* encodes the catalytic subunit of phosphoinositol 3-kinase (PI3K) of the AKT/PI3K/AKT/mTOR pathway, which is essential for protein synthesis, cell growth, proliferation, and survival [54]. Overexpression and constitutive activation of *PIK3CA* leads to inhibition of apoptosis and, thus further tipping the balance toward carcinogenesis [54]. TCGA found 56% of HPV-positive HNSCC showed activating mutations in this gene [53].

Amplification of the *E2F1* was another important HPV-specific finding [53]. *E2F1* encodes a mitogenic transcription factor. E2F1 target genes encode for proteins that are critical for progression of the cell cycle through the G1/S transition [55]. Interestingly, E2F1 is regulated by the Rb tumor suppressor [56], a target of HPV E7 [44]. Overexpression of *E2F1* leads to dysregulation of this cell cycle checkpoint, pushing the cell toward carcinogenesis. TCGA found activating mutations in *E2F1* in 19% of HPV-positive HNSCC, compared to 2% in HPV-negative HNSCC [53].

TCGA found the inactivation of *TRAF3*, found in 22% of HPV-positive tumors, to be another mutation typical of HPV-positive HNSCC [53]. The protein product of *TRAF3* is a negative regulator of the NF-𝜅B signaling, a mediator of cellular proliferation and apoptosis blockade [57]. As an inhibitor of NF-𝜅B, TRAF3 indirectly promotes apoptosis and suppresses proliferation. Inactivating mutations in TRAF3, thus, promote molecular pathways known to plays significant roles in carcinogenesis. Indeed, studies beyond TCGA have identified mutations in TRAF3 and CYLD, another regulator of NF-𝜅B [58], as defining features of a subset of HPV-positive HNSCC [59].

In addition, both HPV-positive and -negative tumors contained recurrent focal amplifications for 3q26/28, a region that involves, among others, the squamous lineage transcription factor SOX2 [53]. This factor controls gene expression in pluripotent stem cells, contributing to the continuation of their pluripotency. Additionally, it plays a role in the reprogramming of somatic cells, reversing their epigenetic configuration back to the pluripotent embryonic state [60]. Evidence of amplification of this gene provides a mechanism for conversion of squamous cells to cancer stem cells, a concept that has been supported by other studies [61].

Other important genes mutated in HPV-positive HNSCC include *DDX3X*, *FGFR2, FGFR3*, *KRAS*, *MLL2/3*, and *NOTCH1* [62] – overlap with mutations in HPV-driven cervical tumors [63].

### Mutational spectrum

Several genome-wide studies including TCGA have found that HNSCC exhibits mutational spectra that appear to be specific to HPV-status [53]. HPV-positive HNSCC display enrichment of cytosine-to-thymidine (C to T) mutations at TpC sites [62,64], which are thought to result from dysregulation of APOBEC. APOBEC comprises a family cytosine deaminases that play key roles in viral immunity by mutating viral DNA and restricting viral replication [65]. However, high levels of APOBEC activity has been observed in several virally-induced cancers to also induce mutation of the host genome, and patterns of APOBEC-related mutagenesis with cytosine deaminase hyperactivity is a common event in virally transformed cancers [62,64].

HPV-related dysfunction of the APOBEC family proteins in HPV-positive HNSCC [62, 64, 66] likely causes non-synonymous mutations within isolated hot-spots. Thus, the prototypical gene mutated by APOBEC dysregulation in HPV-positive HNSCC is *PIK3CA*, in which C to T mutations at two hotspots cause recurring E542K and E545K amino acid substitutions that cause gain of function mutations and result in PIK3CA kinase activation [62,67].

## V. VIRAL INTEGRATION AND CARCINOGENESIS

The insertion of HPV genomic DNA into the host cellular genome, termed viral integration, is not considered a part of the natural life cycle of HPV; however, it may contribute to HPV-related carcinogenesis [68,69]. Viral DNA integration into human genome more often occurs with oncogenic HPV types [70], and is frequently found in HPV-positive OPC [68,69]. Indeed, half of HPV-positive HNSCC (39-71%) have the virus integrated into the host genome [13, 71, 72].

Viral transcripts expressed from integrated viral DNA have been found to be more stable and expressed at higher levels than those from episomal viral DNA [73]. Recent data suggest that integration of oncogenic viruses, including HPV, into the host genome, are non-random [73-76]. A common breakpoint of the viral genome occurs in E2, the virally-encoded transcriptional inhibitor of E6 and E7 (Figure 4) [71,73]. In this manner, integration may result in the upregulation of HPV oncoproteins E6 and E7 [73, 77, 78]. Chromatin regulatory elements near the site of integration also may cause upregulation of integrated E6/E7 genes [72,73]. Such elements include areas of transcriptionally-active “open” chromatin and non-coding transcriptional regulatory sites including enhancers or super-enhancers [79]. Thus, in the case of bovine papillomavirus 1 (BPV1), the correspondence between integration and open chromatin occurs because of BRD4, a reader of acetylated lysine residue 27 on histone 3 (H3K27Ac), mediates tethering of the viral genome to host chromatin [80-82]. Recent data correlate HPV16 integration sites in cervical cancer with super-enhancer domains of open chromatin, enriched by H3K27Ac, MED1 (RNA polymerase II [PolII] transcription subunit), and BRD4 proteins [79]. It is possible that a similar mechanism also mediates viral integration in HPV-positive HNSCC.

**Figure 4:**
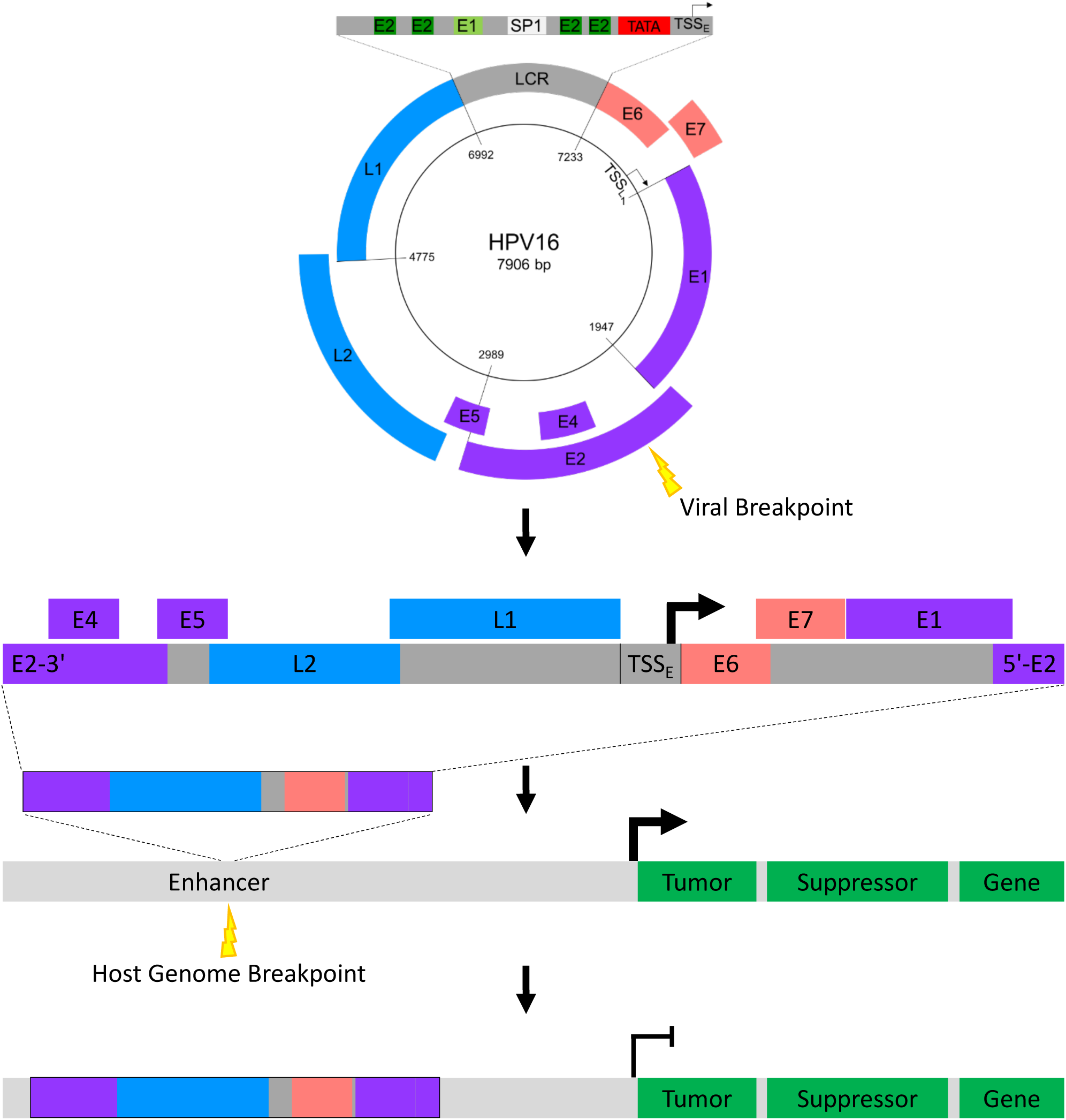
Model for HPV integration promoting tumorigenesis. (A) Breakpoints in the viral genome often occur in the E2 gene. (B) Through interruption of the E2 gene, viral linearization leads to unregulated expression of E6 and E7 viral oncogenes, represented by the bold arrow at TSS_E_. (C) The linearized viral genome inserts into a segment of open chromatin. Although this schematic shows viral DNA integrating at an enhancer, it may also integrate at genic or other transcriptionally active loci. (D) Viral DNA integration may result in the disruption in the expression of nearby genes. In a simplified example depicted here, integration of the viral genome upstream of a host tumor suppressor gene results in its transcriptional inhibition. However, viral integration into gene introns has also been demonstrated gene alter gene expression and alternative splicing [73].

Further, HPV integration promotes genetic instability [68,83], and confers an advantage in cellular proliferation [77]. While some evidence supports a role for HPV DNA integration in HPV-related carcinogenesis, the general principles HPV integration and its mechanistic role in HPV-related carcinogenesis remain controversial.

Early studies employing low-throughput molecular biological approaches sought to elucidate the potential relationship between HPV integration and HNSCC progression. Targeted analysis of viral copy number and RNA expression was conducted on HPV-positive OPC tumor samples by quantitative real time polymerase chain reaction (qRT-PCR) [72]. Twenty-nine of 75 tumor samples (39%) were found to amplify HPV16-human mRNA fusion products, evidence consistent with viral integration. Comparison of HPV-integrated and episomal OPC samples found no difference in viral copy number, expression of HPV16 interrupted genes, or expression of viral genes E2, E6, and E7 between HPV-integrated and episomal tumors [72]. This data suggested that HPV16 integration had no impact on HPV-dysregulated human transcripts and or the expression of key viral oncogenes [72]. In a subsequent qRT-PCR-based analysis of 7 HPV-positive HNSCC cell lines with integrated and/or episomal viral DNA, Olthof et al drew similar conclusions [71]. This latter study showed that viral E2, E6 and E7 gene expression was independent of viral load and the number of viral integration sites, suggesting that integration did not affect viral gene expression [71].

A more recent analysis of high-throughput data from TCGA, however, partially contradicted these findings [71,72]. Parfenov et al defined HPV status by interrogating 279 HNSCC samples with associated RNA-sequencing data for samples expressing HPV E6 transcripts [73]. Viral integration status was then analyzed from associated DNA-sequencing data in the HPV-positive samples. Of 35 HPV-positive HNSCC samples, each of 25 tumors exhibited non-random viral integration into 1 to 16 regions of the human genome, leaving *E6* or *E7* intact in all but one case. Interestingly, seventy-five of the 103 total integration events occurred in or within 20 kilobases of a known gene. Higher detail insertion-site analysis identified several putative mechanisms demonstrating that non-random HPV integration may confer a selective advantage to tumor cells and promote tumor progression.

HPV was found to integrate into and alter the function of several known tumor-associated genes, including *RAD51B, ETS2, PDL1.* One tumor harbored three viral insertions into intron 8 of the *RAD51B* gene. Expression analysis of this tumor demonstrated significantly elevated expression of exons 9-11 and 13 and alternative transcripts unlikely to produce functional RAD51B protein. RAD51B is an essential component of DNA double-strand break repair machinery and, consistent with its role as a tumor suppressor gene, loss-of function *RAD51B* mutations have been reported in cancer [84]. Monoallelic HPV integration was detected in *ETS2*, another tumor suppressor, resulting in significant reductions in the expression of exons 7 and 8. This suggests alternative splicing pattern potentially may lead to biallelic loss of *ETS2* function by a dominant negative effect [85]. Similarly, intronic HPV integration into the *PDL1* gene resulted in augmented expression of an alternative *PDL1* transcripts. Alternative PDL1 transcripts have been associated with poor prognosis renal cell carcinoma [86].

Parfenov et al also identified HPV integration in intergenic loci known to regulate tumor-associated genes *TPRG1, TP63,* and *KLF5* [73]. The concomitant transcriptional upregulation of these target genes was directionally consistent with tumor-promoting roles of these genes [13,87]. Similarly, HPV integration upstream of the *NR4A2* oncogene associated with *NR4A2* overexpression [88]. Consistent with these findings, analysis of 8 HPV-positive HNSCC cell lines by qRT-PCR identified HPV integration events and evidence consistent with dysregulation of several cancer-related genes *TP63, DCC, JAK1, TERT, ATR, ETV6, PGR, PTPRN2,* and *TMEM237* [89]. Although preliminary, these findings set forth putative mechanisms by which HPV integration may promote tumor progression.

## VI. TRANSCRIPTOME-LEVEL ALTERATIONS OF HPV-RELATED HNSCC

### Gene Expression Signatures and HNSCC Subtypes

Not until recently have comprehensive studies of gene expression in HPV-positive and HPV-negative HNSCC tumor types been conducted to distinguish them based on underlying genetic aberrations. In addition to further distinguishing the molecular features of HPV-positive HNSCC, this research has led to the proposal of HNSCC subtypes that better capture its biological heterogeneity and may facilitate the development of subtype-specific targeted therapeutics.

This line of study was pioneered by Seiwert et al, who found that HPV-positive tumors showed a distinct genetic profile with unique mutations in *DDX3X, CYLD*, and *FGFR,* as well as enrichment for PI3K pathway alterations and rarer *KRAS* mutations common for HPV-negative HNSCC tissues. As an opposite, the mutational profile in HPV-negative HNSCC included enrichment for mutations in *TP53, CDKN2A, MLL2, CUL3, NSD1,* and *PIK3CA* genes, and copy-number increases in *EGFR*, *CCND1*, and *FGFR1*. The group also found that somatic aberrations in DNA-repair genes (*BRCA1/2*, Fanconi anemia genes, and *ATM*) may contribute to chemo- and/or radiosensitivity of HPV-positive tumors. These DNA-repair genes mutations occurred in HPV-positive tumors in non-/light smokers [66]. Considering a recent report that *RAD51B*, a BRCA2 cofactor [90], was a potential integration target for HPV16 leading to loss of its DNA-repair function, may indicate that the role of DNA repair gene disruption in HPV-positive HNSCC carcinogenesis warrants further investigation.

One group, Keck et al, established subtype classification using gene expression-based consensus clustering, copy number profiling, and HPV status on a cohort of 134 locoregionally advanced HNSCCs. They identified five subtypes of HNSCC, including, most importantly, two biologically distinct HPV subtypes. The first HPV-positive HNSCC subtype is the classic HPV positive subtype (CL-HPV). This subtype is characterized by significant enrichment of the polyamine degradation pathway, which is relevant for detoxification, for example, related to tobacco use. More than 40% of CL-HPV tumors show keratinization, and this subtype has a higher proliferation rate compared to the other subtypes. The second HPV-positive subtype is the inflamed/mesenchymal HPV-positive subtype (IMS-HPV), which shows an immune and mesenchymal phenotype. Prominent tumor infiltration with CD8+ lymphocytes is characteristic of IMS-HPV, independent of HPV status. This tumor subtype shows no keratinization, and exhibits elevated mesenchymal markers. Establishment of these subtypes carries translational implications for the development of subtype-specific biomarkers, and it provides a biological justification for the use of different treatment approaches among HPV-positive HNSCC based on subtype [91].

### The emerging role of ErbB family proteins in HPV-positive HNSCC therapeutics

ErbB family proteins are encoded by 4 genes that play a central role in carcinogenesis: *ERBB1* (also known as *HER1* or *EGFR*), *ERBB2* (also known as *HER2* or *CD340*), *ERBB3* (also known as *HER3*), and *ERBB4* (also known as *HER4*) [92,93]. Pollock et al published a paper in 2015 which reported that expression of total HER2, total HER3, HER2:HER3 heterodimers, and the HER3:PI3K complex were significantly elevated in HPV-positive HNSCC [94]. They also reported that Afatinib, an anti-ErbB family small molecule inhibitor, significantly inhibited cell growth compared to cetuximab in HPV-positive cetuximab-resistant HNSCC cell lines. These findings suggested that targeting ErbB family receptor tyrosine kinases may be a potentially effective treatment for HPV-positive HNSCC.

However, these results were not supported by the LUX Head and Neck 1 trial [95,96], which showed that both patients with p16-positive tumors and patients whose tumors progressed after cetuximab therapy derived less benefit from afatinib. One possible explanation for this apparent discrepancy was a difference in definitions; for example, the LUX study considered previous exposure to cetuximab equivalent to cetuximab-resistance [95,96].

The discrepancy was addressed with a recent study investigating the relationship between HPV oncoproteins and HER3-mediated signaling, and the role of HER3 as a potential molecular target in HPV-positive HNSCC [97]. This study proposed a novel mechanism by which HPV can promote tumor growth through the mTOR/AKT/PI3K pathway. HER3 expression was found to be significantly increased in HPV-positive cell lines compared to HPV-negative cell lines. Brand et al then investigated the role of E6 and E7 oncoproteins in *HER3* expression. They found that E6/E7 gene silencing led to diminished *HER3* production, while increased E6/E7 oncoprotein production led to increased *HER3* production. The results indicate a relationship between HPV infection and *HER3* expression that is mediated by the E6 and E7 oncoproteins [97].

Having found evidence to support this relationship, the group then studied how *HER3* expression might affect cellular proliferation and tumor growth in HPV-positive HNSCC cell lines. Their suspicions were confirmed; genetic ablation of *HER3* resulted in statistically significant inhibition of cellular proliferation (21-55%) [97]. Furthermore, *HER3* knockdown led to reduced AKT and RPS6 phosphorylation [97]. The reduction of these proteins, which are components of the same mTOR/AKT/PI3K pathway discussed in the genomics section of this review, suggests that HER3 exerts its proliferative effect by signaling through the mTOR/AKT/PI3K signaling pathway [97].

Based on their findings suggesting that *HER3* is overexpressed in HPV-positive cell lines, patient-derived xenografts, and human tumors, Brand et al investigated the sensitivity of HPV-positive preclinical models to anti-HER3 monoclonal antibody therapy, such as KTN3379. KTN3379 was shown to be effective in growth inhibition in 4 out of 4 HPV-positive HNSCC cell lines, versus 2 out of 5 HPV- lines [97]. KTN3379 inhibited phosphorylation of HER3 in all HNSCC cell lines regardless of HPV status, but analysis of downstream PI3K signaling revealed that phospho-AKT and phospho-RPS6 were more significantly inhibited in three HPV-positive cell lines versus three HPV-negative cell lines [97]. HPV-positive human HNSCC tumors xenografted into mice also showed statistically-significant reduction in tumor volume, as well as reduced levels of total *HER3*, phospho-HER3, phospho-AKT, and phospho-rpS6 [97]. These findings indicate that anti-HER3 therapy may have potential as an tumor type-specific HPV-positive HNSCC therapeutic target. Further supporting a potential role of ErbB family proteins, retrospective study of a patient with HPV-positive OPC with complete response to HER2-targeted therapy found that a dermal metastasis from this patient exhibited HER amplification[98].

## VII. EPIGENETIC ALTERATIONS IN HPV-RELATED HNSCC

Additional amplification of the host genome at the site of HPV integration [53], and dysregulation of p53 and Rb tumor-suppressor genes by HPV oncoproteins E6 and E7 do not fully account for the genome-wide spectrum of gene expression changes in HPV-positive HNSCC [53]. Chromatin structure and epigenetic modifications, such as DNA methylation and histone-modifications, regulate gene expression, and their changes may explain genome-wide dysregulation found in HPV-positive HNSCC.

### Methylation

The drive to understand the molecular underpinnings of HPV-positive HNSCC and, based on that knowledge, to develop biomarkers predictive of prognosis and treatment has now progressed to include epigenetic changes. The disruption of epigenetic signatures can cause alterations in gene function that lead to malignant changes in cells [99]. DNA methylation, the addition of a methyl group to a DNA molecule, especially the C5 position of cytosine, is the most common form of epigenetic modification. Given how often promoter methylation is found to be the mechanism of action in transcriptional silencing in HNSCC [100], an exploration of the unique methylation patterns inherent in HPV-positive HNSCC may be helpful in creating clearer molecular profiles and tailored treatment plans.

One such exploration was conducted by Kostareli at al, found a negative correlation between gene promoter hypermethylation and gene transcript levels in *ALDH1A2^lo^*, *OSR2^lo^*, *GATA4^hi^*, *GRIA4^hi^*, and *IRX4^hi^* in HPV-related tumors. These hypermethylation signatures were positively correlated with better patient prognostic outcomes. These findings may serve as a basis for the accurate identification of patients who are at greater risk of treatment failure, and distinguish them from patients with better prognostic indicators. Patients at a higher risk of treatment failure would benefit from multimodal treatment options, whereas patients with more favorable prognosis may benefit from lower toxicities associated with treatment de-escalation [101].

Sartor et al HPV-positive HNSCC overall were found to have greater DNA methylation, both in terms of targeted genic methylation and global methylation, compared to HPV-negative HNSCC. The group found higher promoter methylation of polycomb repressive complex 2 target genes and increased expression of *DNMT3A,* in HPV-positive cells, implicating this DNA methyltransferase in the methylation patterns observed in HPV-positive HNSCC. *RUNX2, IRS-1* and *CCNA1* had higher methylation and lower expression in HPV-positive cells, whereas *CDKN2A* (the surrogate marker of HPV infection [48]) and *KRT8* had higher methylation and lower expression in HPV-negative cells [102].

Similar to Sartor et al, the independent group of Lleras et al reported a greater level of differential DNA methylation in HPV-positive OPC cases compared to HPV-negative cases, and identified that this increase was due in large part to hypermethylation of CpG islands in HPV-positive cases. There were 28 genes found in association with these methylation patterns in HPV-positive OPC. Some novel findings included the association of differential methylation patterns with *ALX4*, *CUTL2*, and *HOXA7*, as well as previously identified cancer-related genes such as *FBX039*, *IGSF4*, *PLOD2* and *SLITRK3*. Two genes identified in this study which were also reported in the Sartor group discussed above included: *CDKN2A*, and *HOXA7*. Notably, Lleras et al found increased promoter hypermethylation in HPV-positive cases, whereas Sartor found increased promoter hypermethylation in HPV-negative cases. A possible explanation for this discrepancy may be found in study design: Sartor et al excluded samples containing less than 40% tumor from subsequent analysis for nodal metastasis signature and HPV status, while Lleras et al included such samples in the initial identification of hyper- and hypo-methylated CpG loci [103].

### Chromatin

Out of different functional chromatin domains, promoters and enhancers play a major role in gene expression regulation via recruitment of specific transcription factors (TFs) to DNA [104]. The enhancers may drive transcriptional dysregulation responsible for the development and progression of HPV-positive HNSCC at a genome-wide scale. [78, 104-110]. HPV-E6/E7 proteins were also found to interact with p300 and MYC, main constituents of enhancer regions [111-114].

These data suggest that targets of viral oncoproteins play the central role in activation of super-enhancers, the epigenetic regulators of gene expression.

Current literature suggests that super-enhancers at the site of HPV integration upregulate the expression of integrated HPV-E6/E7 oncogenes [79]. These overexpressed viral oncoproteins most likely activate their direct or indirect protein targets, which activate super-enhancer complexes genome-wide in disease-specific manner [115]. These disease-specific super-enhancers with unique components can be therapeutically targeted [115]. Indeed, the leading HNSCC oncoproteins NF-κB and STAT [116], were also found to participate in enhancer formation during Epstein Barr Virus-caused carcinogenesis [115] In addition, since E6/E7 lack enzymatic activities and don’t directly bind to DNA they exert their functions by binding with a vast number of cellular proteins [117]. Published data suggest that direct or indirect targets of HPV E6/E7 affect the chromatin landscape of cancer cells [118-120]. Indeed, E7 induces the expression of KDM6B (demethylase of H3K27me3, the mark of repressive chromatin), which increases chromatin accessibility and enhances gene transcription [118]. HPV16 E7 activates HATs (histone acetyltransferases) and inactivates HDACs (histone deacetylases) [119,120], leading to acetylation of histone tails and chromatin opening. Additionally, binding and inactivation of E2F6, Rb, and p130 by HPV E7 induces acetylation of H3K9 and methylation of H3K4, both of which are marks of open chromatin and active transcription [119, 121, 122]. These and other data suggest that reorganization and activation of the chromatin are primary mechanisms of HPV-related carcinogenesis.

## VIII. CONCLUDING REMARKS

In this review, we briefly described the prevailing molecular alterations observed in HPV-positive HNSCC, a disease currently lacking effective targeted therapeutics. The current literature suggests that viral oncoproteins lead to significant epigenetic cascades and that aberrant chromatin structure plays an important role in the integration of HPV into the host genome. A “chicken and the egg” question remains unanswered for the potentially interdependent phenomena of HPV integration and HNSCC aberrations in chromatin structure, and requires further mechanistic investigation. Furthermore, unlike cervical cancer in which most tumors exhibit viral integration, HPV-positive OPC often display episomal HPV DNA. Whether these observations represent a divergence in the mechanisms of HPV-related carcinogenesis in HNSCC also remains an unanswered question. Epigenetic dysregulation and APOBEC-signature mutations most likely drive genome wide dysregulation of the gene expression. Many of these genes, including ErbB-family proteins, represent promising targets for the development of tumor type-specific therapeutics. The significant progress in the understanding of HPV-related carcinogenesis in HNSCC reviewed here will serve as the foundations toward the development of novel diagnostic and therapeutic modalities to quell the burden imposed by this emerging viral malignancy.

## CONFLICTS OF INTERESTS

The authors have no conflicts of interest to disclose.

## Acknowledgments

D.A.G. receives funding from NIDCR, grant number R21-DE025398.

